# Exploring the properties of the left angular gyrus using TMS-evoked potentials

**DOI:** 10.1101/2022.11.15.516568

**Authors:** Dominika Sulcova, Yasmine Salman, Adrian Ivanoiu, André Mouraux

## Abstract

The angular gyrus (AG) is involved in multiple cognitive processes and its structural alterations are commonly observed in many neuropsychiatric syndromes. Since changes in excitability may precede structural changes and clinical symptoms, there is a need for diagnostic tools assessing the functional state of hub brain regions like the AG. The combination of transcranial magnetic stimulation (TMS) and electroencephalography (EEG) can provide such functional readouts by probing the brain response to direct stimulation.

This study aimed to characterize TMS-evoked potentials (TEP) elicited by AG stimulation, determine optimal stimulation parameters, and identify TEP biomarkers of AG function. We recorded AG-TEPs in 19 subjects using four TMS orientations and three intensities and compared TEP spatiotemporal features using topographic dissimilarity and microstate analyses. Additionally, we explored the relationship between AG-TEP topography and TMS-evoked muscular activity.

Our results showed topographic variability in AG-TEP components P25, N45, and N75. The P25 topography was sensitive to TMS orientation, while the topography of N45 and N75 was highly dependent on both coil orientation and intensity. Interestingly, we found that TMS-evoked muscular activity was also dependent on both these parameters and strongly related to the occurrence of a specific topographic pattern, which therefore possibly corresponds to the somatosensory brain response to muscle contraction.

We conclude that the early AG-TEP component P25 likely reflects neural processes triggered by direct AG activation and could provide an index of local excitability. Later components N45 and N75 must be interpreted with caution as they may primarily reflect peripherally evoked activity.

## Introduction

Located right at the junction of temporal, occipital and parietal lobes, the angular gyrus (AG) represents a highly connected cortical hub (Hagmann et al., 2008; Kernbach et al., 2018; Tomasi & Volkow, 2011) that is implicated in a vast range of cognitive processes (for an exhaustive review see (Seghier, 2013)). Amongst others, it constitutes one of the core nodes of the default mode network (DMN), which is a collection of large-scale subnetworks mediating internally driven mental processes (Buckner & DiNicola, 2019; Greicius, Krasnow, Reiss, & Menon, 2003; Shehzad et al., 2009). Due to its widespread involvement, any damage or functional impairment of the AG inevitably impacts cognitive performance (Ardila, 2014). Correspondingly, structural and connectivity abnormalities of the AG have been reported in a variety of neurologic and psychiatric disorders including Alzheimer’s disease (Berron, van Westen, Ossenkoppele, Strandberg, & Hansson, 2020; Jagust et al., 2006; Smith et al., 2007), schizophrenia (Chahine, Richter, Wolter, Goya-Maldonado, & Gruber, 2017; Torrey, 2007), amyotrophic lateral sclerosis (Agosta et al., 2013; Sakurai et al., 2021), and Huntington’s disease (Macdonald, Halliday, Trent, & McCusker, 1997). As it is likely that these pathologic alterations are preceded by more subtle changes in local cortical excitability or connectivity, there is a strong incentive to explore functional properties of the AG, particularly in early-stage patients and preclinical populations.

Most previous studies used functional magnetic resonance imaging (fMRI) to assess the AG in the context of large-scale brain networks. This technique approximates the activity of target brain regions by measuring the hemodynamic brain response (Attwell & Iadecola, 2002; Ogawa, Lee, Kay, & Tank, 1990), so the signal only shows relatively slow fluctuations of cortical activation (Cole, Smith, & Beckmann, 2010). By consequence, fMRI does not allow to capture the underlying neural dynamics occurring at the millisecond timescale that could provide specific information about the local excitability or the causal connectivity within the stimulated brain network (Bressler & Tognoli, 2006).

It is now possible to obtain such measures by combining the spatial precision of transcranial magnetic stimulation (TMS) with the high temporal resolution of electroencephalography (EEG). While TMS engages neuronal circuits in a well-defined cortical region (Barker, Jalinous, & Freeston, 1985; Hallett, 2007), EEG records the activity evoked by the TMS perturbation and its subsequent spread through the connected brain areas (Bergmann, Karabanov, Hartwigsen, Thielscher, & Siebner, 2016; Ilmoniemi & Kičić, 2010). The magnitude and spatiotemporal distribution of TEPs can be thus used to evaluate the excitability of the stimulated cortex (Bergmann et al., 2016; Ferreri & Rossini, 2013; Mäki & Ilmoniemi, 2010) and gain insight into the causal connectivity within corresponding brain networks (Bergmann & Hartwigsen, 2021; Bortoletto, Veniero, Thut, & Miniussi, 2015; Hallett et al., 2017; Rogasch & Fitzgerald, 2013).

To this date, only few TMS-EEG studies have targeted the AG and the experimental settings they used, including the exact location of the site of stimulation, varied from one study to another (Freedberg, Reeves, Hussain, Zaghloul, & Wassermann, 2020; Lauro et al., 2014; Ozdemir et al., 2020; Ozdemir et al., 2021; Ross et al., 2022; Šulcová, Salatino, Ivanoiu, & Mouraux, 2022). Nevertheless, the available data indicate that it is possible to record AG-TEPs that are stable across sessions and have spatiotemporal profiles specific for individual subjects (Ozdemir et al., 2021). Furthermore, EEG source reconstruction revealed a target-specific propagation of activity from the stimulated AG across different nodes of the DMN, thereby hinting at the utility of AG-TEPs for the evaluation of this resting-state network (Ozdemir et al., 2020). However, to establish AG-TEPs as a clinically relevant tool for the assessment of local cortical excitability of the AG as well as the perturbational connectivity in the DMN, it is crucial to characterize the physiological profile of AG-TEPs, to identify the components that genuinely reflect the activity within the stimulated network, and to find the optimal combination of stimulation parameters.

It is generally agreed that the stimulation intensity as well as the orientation and direction of the magnetic field in relation to the target cortical area determine which neuronal populations are stimulated by TMS and how efficiently (Rossini et al., 2015; Siebner et al., 2022). Because neuronal cells seem to be most susceptible to electric currents flowing in the direction from the dendrites to the axon, cortical columns in sulcal walls should be most activated when the magnetic field is oriented perpendicularly to the axis of the targeted gyrus (Fox et al., 2004). However, the horseshoe shape of the AG offers several possible ways to place the TMS coil, each orientation presumably prioritizing different parts of the sulcal wall of the area. This is a particularly relevant notion when we consider that the AG is not a homogenous region but rather consists of several subdivisions that show distinct cytoarchitecture (Caspers et al., 2008; Caspers et al., 2013), connectivity profile (Mars et al., 2011) and function (Nelson et al., 2010; Numssen, Bzdok, & Hartwigsen, 2021; Seghier, Fagan, & Price, 2010). Therefore, it can be expected that different combinations of stimulation parameters may activate the target area with variable efficiency and/or preferentially engage different brain circuits. Moreover, stimulation parameters such as coil placement may also differentially influence undesired activation of peripheral nerves and muscles (Conde et al., 2019; Mutanen, Mäki, & Ilmoniemi, 2013).

In the present study, we aimed to describe the spatiotemporal profile of TEPs evoked by stimulation of the left AG, to assess the impact of different stimulation parameters on the recorded responses and, if possible, to identify TEP components reflecting the activity directly evoked within the AG. To this end, we applied TMS to the posterior-ventral part of the AG, an area that previously showed strong connectivity to the hippocampus and other nodes of the DMN (Mars et al., 2011; Uddin et al., 2010). AG-TEPs were recorded using four different stimulation orientations (defined by coil placement and current direction) and three stimulation intensities and compared across conditions using the analysis of topographic dissimilarity and the microstate analysis (Koenig, Stein, Grieder, & Kottlow, 2014; Šulcová et al., 2022). Based on the assumption that two different topographies represent the activity of two distinct sets of cortical generators (Michel et al., 2004), these two methods allow making inferences about changes in underlying TEP sources without estimating their exact location. Finally, we explored the association between the magnitude of TMS-evoked contraction of head muscles and prominent topographic patterns of AG-TEPs identified by the topographic analysis.

## Material and Methods

### Participants

19 healthy volunteers participated in the study (9 males, mean age ± SD: 23.78 ± 3.34 years). Before enrolling, candidates filled a questionnaire to control for contraindication to experimental procedures (Rossi, Hallett, Rossini, Pascual-Leone, & Group, 2009). Accepted subjects gave a written informed consent and were financially compensated for their participation. All performed procedures were conducted according to the Declaration of Helsinki and approved by the Ethical Committee of Catholic University of Louvain and University Clinics Saint-Luc.

### Experimental design

TEPs were recorded from the AG using different stimulation orientations defined by the position of the TMS coil - across or along the superior temporal sulcus (STS) – and the direction of the electric current in the TMS coil - normal or reversed phases (Fig. 1c). For each of the resulting four combinations, we tested three stimulation intensities based on the individual threshold to evoke motor response at rest (resting motor threshold, rMT) – 100, 120 or 140% rMT. The experiments were conducted in two sessions (approximately 3.5 h) separated by one week, each testing one of the two coil positions (Fig. 1d).

**Fig. 1.**
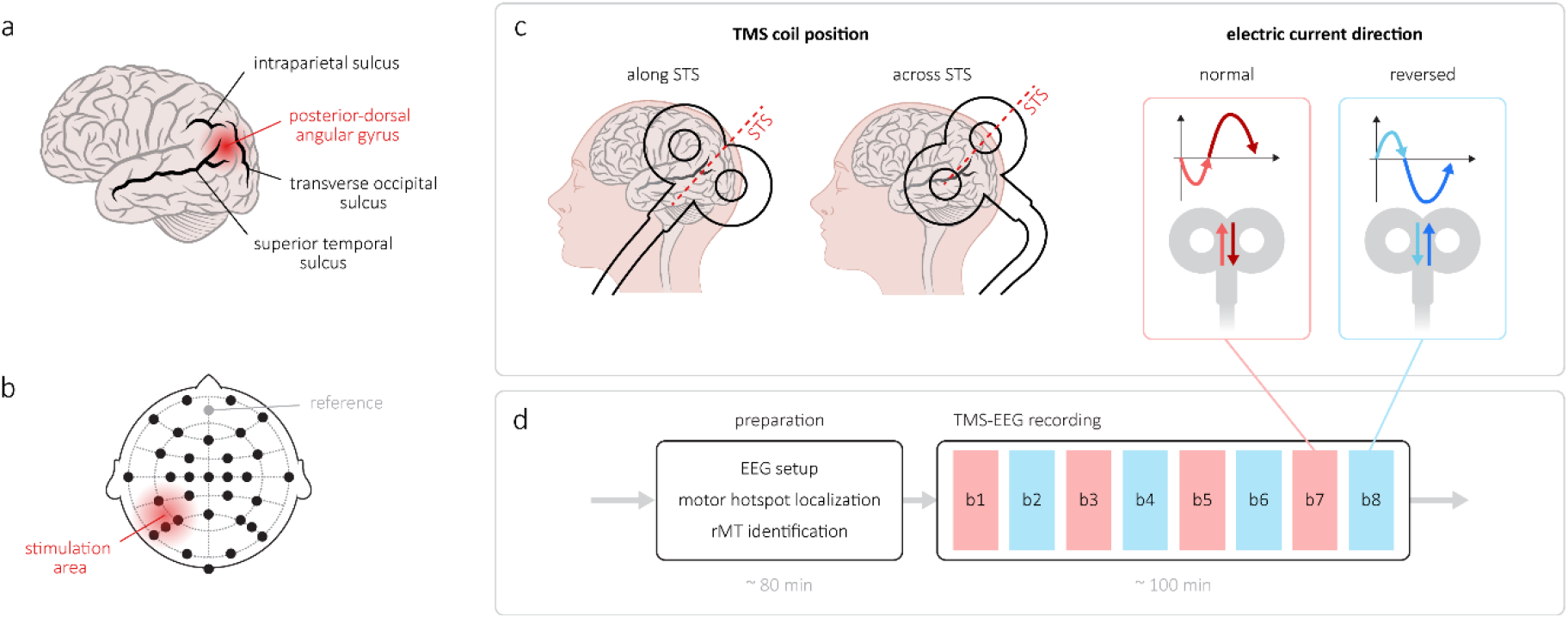
Experimental design. **(a)** The target site was located in the posterior-dorsal AG on the gyral crown defined relative to the position of the most prominent sulci marked in the figure. **(b)** EEG layout used in the study comprised 32 active electrodes (black), the reference is shown in grey, the red shading approximates the area under the TMS coil. **(c)** Four tested stimulation orientations were defined as the combination of two different TMS coil placements – along and across the axis of the superior temporal sulcus (STS; marked in red) and two directions of the electric current in the coil – normal (prominent second phase in dark red) and reversed (prominent phase in dark blue). **(d)** One experimental session contained 8 blocks of TMS-EEG (each approximately 6.5 mins long) with the coil placement constant but the current in the coil alternating between the blocks.

### EEG recording

The EEG was recorded with the NeurOne EEG system (Bittium NeurOne Tesla; Bittium Corporation, Oulu, Finland) using a 32 channel EEG cap mounted with TMS-compatible Ag/AgCl electrodes (EasyCap GmbH, Herrsching, Germany) (Fig. 1b). The signal was referenced to FCz and recorded at 20 kHz sampling rate with 5000 Hz low-pass filter and DC filter, with electrode impedances kept below 5kΩ. Subjects were seated in a comfortable chair and fixing their gaze to a stable point. A layer of thin plastic and another textile cap were placed over the EEG cap to minimize EEG artifacts caused by the contact of electrodes with the TMS coil and subjects were listening to a custom masking noise (Russo et al., 2022) to minimize auditory artifacts.

### TMS stimulation

A 3D T1-weighted structural MRI of the head was acquired in advance (1×1×1 mm; 3 T Achieva; Philips Healthcare, Amsterdam, The Netherlands) and 3D models of the brain surface and the scalp were reconstructed in the Visor2 neuronavigation system (Visor 2.3.3; Advanced Neuro Technologies, Enschede, The Netherlands). The posterior-ventral AG was located on the brain model dorsally to the ascending branch of the STS (Fig. 1a), its position was projected on the scalp model and two TMS coil targets were created, one with the axis along the STS and the other perpendicular to it. The model was co-registered with subject’s head using landmarks (nasion and tragi) and head-shape matching (Gugino et al., 2001). The position of the head and the TMS coil was monitored with a Polaris optical tracking system (Polaris Spectra; Northern Digital Inc. Europe, Radolfzell, Germany) and the neuronavigation ensured the accurate coil placement throughout the stimulation.

Biphasic TMS pulses were delivered manually using a MagPro stimulator (MagPro X100 with MagOption; MagVenture, Farum, Denmark) with a figure-of-eight coil (75 mm; MagVenture, Farum, Denmark) placed tangentially to the scalp and aligned with the target. The intensity of stimulation was determined based on the rMT, which was identified as the minimal intensity eliciting a motor response with an amplitude ≥ 50 μV in at least 5/10 trials (Rossini et al., 2015). The average (±SD) rMT corresponded to 44.3 ± 8.5% of the maximum stimulator output (MSO). Each experimental session contained 8 blocks of 75 TMS pulses (jittered interstimulus interval 4-6 s) that were randomly assigned one of the tested stimulation intensities (Fig. 1d), delivering 100 stimuli in total for every condition.

### Data pre-processing

EEG recordings were pre-processed offline using Matlab (MathWorks, Inc., Natick, Massachusetts, United States), Letswave 6 (an open-source EEG/EMG signal processing toolbox, https://www.letswave.org/) and custom scripts following the pipeline introduced by Rogasch (Rogasch et al., 2014).

The signal was re-referenced to the common average, epoched (-1000 to 2000 ms relative TMS), and the DC shift and linear trend were removed. The large muscle artifact was replaced by cubic interpolation (-5 to +10 ms) and the signal was downsampled to 2 kHz. A first round of Independent Component Analysis (ICA, FastICA algorithm (Hyvärinen & Oja, 2000)) was performed to remove components containing the sharp leftover tail of the muscle artifact. Data were bandpass filtered (0.1-80 Hz, Butterworth, 4^th^ order) and notch filtered (50 Hz, FFT linear filter, width 2 Hz, slope 2 Hz). All epochs were visually inspected to discard those containing excessive noise, the average final number of epochs per condition was 83.7 ± 7.9.

A second round of ICA was used to remove remaining artifacts, such as eye movements, tonic muscle activity and electrode noise, as well as the remains of the muscular artifact. Suspicious components were identified with the Multiple Artifact Rejection Algorithm (MARA) (Winkler, Haufe, & Tangermann, 2011) and evaluated visually based on their topography, time course and frequency content. For more detailed information on this step, the reader may consult the supplementary material of (Dominika Sulcova, Adriana Salatino, Adrian Ivanoiu, & Andre Mouraux, 2022a).

Finally, the TEPs were baseline corrected (-200 to -5 ms) and group averaged.

### Data analysis and statistics

AG-TEPs were evaluated between 10 and 300 ms post stimulus. The peaks within this window were identified in the group average waveforms separately for each stimulation orientation (intensities pooled) and labelled according to their prominent polarity (P = positive, N= negative) and approximate peak latency (ms). Latencies were extracted from local maxima of the Global Field Power (GFP; Equation 1) (Lehmann & Skrandies, 1980).

Topographic analysis was performed in Ragu (Matlab toolbox for randomization-based analysis of EEG event-related potentials (Koenig, Kottlow, Stein, & Melie-García, 2011)) and using custom Matlab scripts. First, outliers were identified based on the Mahalanobis distance between subjects (Wilks, 1963) and excluded. The subsequent steps followed the pipeline indicated in (Habermann, Weusmann, Stein, & Koenig, 2018), always using 5000 permutations for the randomization and the level of significance set to 0.05. Here, we provide a brief outline of the process and invite the reader to see (Dominika Sulcova, Adriana Salatino, Adrian Ivanoiu, & André Mouraux, 2022b) for a detailed description.

Because topographic analysis operates with normalized data, it disregards potential differences in TEP amplitude. Therefore, we additionally evaluated the influence of stimulation parameters on the magnitude of AG-TEP components and these results are reported in the Supplementary materials.

#### Planned comparisons

The effect of two factors on AG-TEPs as well as their interaction were evaluated: (1) *TMS orientation*, with four levels corresponding to different combinations of TMS coil position and electric current direction (*along-normal, along-reversed, across-normal*, and *across-reversed*); (2) *TMS intensity*, with three levels (100, 120, and 140% rMT).

#### Test for topographic consistency (TCT)

TCT was conducted separately for each tested condition to identify time intervals showing significant topographic homogeneity across individuals. The GFP of the group average TEP was used as a quantifier and tested against an empirically obtained distribution computed from signals spatially scrambled at subject level.

#### Analysis of topographic similarity

TEP topographies were compared using the measure of Global Dissimilarity (DISS; Equation 2) introduced by (Lehmann & Skrandies, 1980). For each planned comparison, DISS was calculated based on group average TEPs (normalized at subject level by GFP) and a point-by-point statistical randomization test (TANOVA) was conducted to determine the time intervals of significant difference. The clusters of significant p values were corrected for multiple comparisons by applying a Ragu procedure called “Global Duration Statistics”, which determines the minimum necessary number of consecutive significant time points based on the data obtained by randomization. For each factor, we calculated the momentary percentage of total explained variance (%EV), which allowed us to identify temporal maxima of topographic dissimilarity. Mean TEP topographies were extracted at these latencies and post-hoc comparisons were performed using channel-wise paired t-tests. Obtained t values were plotted as t-maps to describe the distribution and magnitude of topographic differences. The significance level was Bonferroni corrected and only significant comparisons were reported.

#### Microstate analysis

Microstate analysis was used to evaluate temporal properties of observed topographic differences. First, the optimal number *n* of microstate classes was determined using the cross-validation method (Koenig et al., 2014). In the next step, *n* microstate classes were identified using the k-means clustering algorithm (Murray, Brunet, & Michel, 2008) with 250 restarts. Each timepoint within each group average TEP was then labelled with the class map that yielded the highest spatial correlation with the momentary topography, thereby segmenting the waveform into intervals of dominance of one of the *n* microstate classes. We then compared temporal properties (*onset, offset*, and *mean duration*) of each class across datasets. For each comparison and microstate, we quantified the observed effect by computing its variance across levels and tested it against an empirically obtained distribution of values true under the null hypothesis.

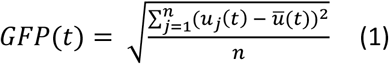

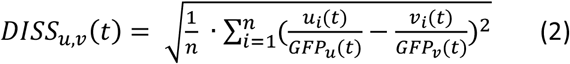

##### Equations of (1) Global Field Power GFP and (2) Global Dissimilarity DISS

Calculated at each timepoint *t, n* is the number of electrodes, *u*_*i*_ and *v*_*i*_ represent the voltage measured at one electrode in two different conditions, while *ū* is the mean voltage in a single condition.

## Results

In the text, we will refer to TEP datasets by the combination of stimulation parameters used: *orientation (along/across) – current (normal/reversed) – intensity (100/120/140)*.

### Spatiotemporal characteristics of AG-TEPs

One individual was identified as outlier and excluded (final n = 18) and remaining datasets were tested for topographic consistency, which was confirmed for the whole duration of the analysed time window (10 – 300 ms post stimulus) with the exception of five short intervals: 34 – 51 ms in *along-reversed-120*, 12 – 19 and 32.5 – 39 ms in *along-reversed-140*, 50.5 – 61 ms in *across-normal-100* and 10 – 14.5 ms in *across-normal-140*. These epochs were not removed from subsequent analysis but were considered when interpreting the results.

AG-TEPs of all conditions comprised six components that were distinguishable already at the individual subject level. To describe their general properties, data from the three stimulation intensities were pooled together for each stimulation orientation (Fig. 2). Peak latencies of these components as identified in the group average GFP were similar across all four datasets, whereas their associated topographies visibly differed in the three earlier peaks (Fig. 2, upper rows). For future reference, AG-TEP components were labelled consistently in all conditions: P25, N45, N75, N100, P200 and P280.

**Fig. 2.**
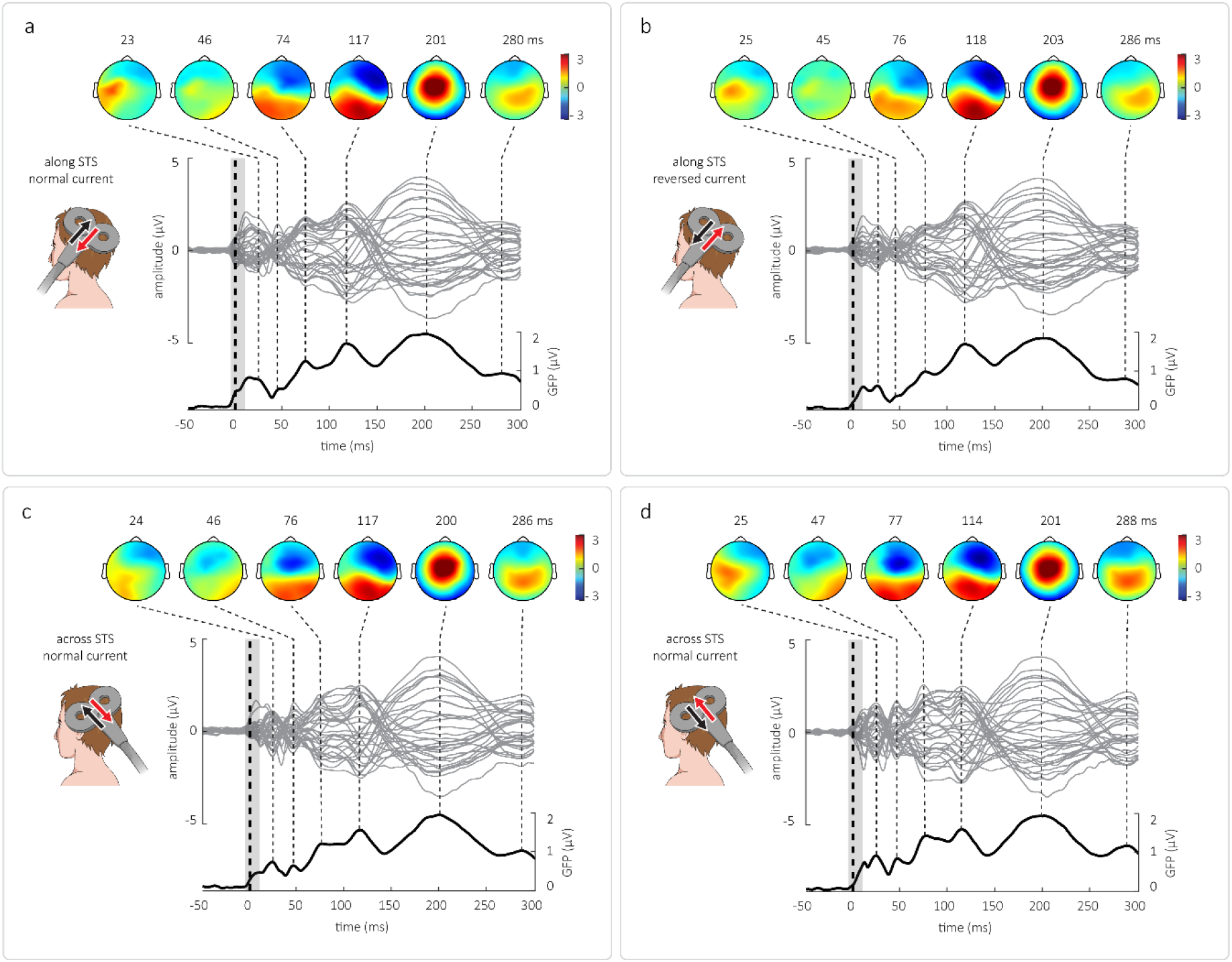
Spatiotemporal profile of AG TEPs obtained with the TMS coil placed along (a, b) or across (c, d) STS and the current flowing in the normal (a, c) or reversed (b, d) direction. For the purposes of visualization, TEPs evoked by all three intensities were averaged. The current direction in the coil is marked by a black (phase 1) and a red (phase 2) arrow. Grey lines in the middle section represent the signal of all recorded EEG channels, the TMS stimulus is denoted by a thick dashed line and the interpolated interval is shaded in light grey. The GFP of the average signal is plotted at the bottom and identified TEP components are marked by dotted lines that are associated with peak topographies displayed in the upper section; peak latencies in ms post stimulus are noted above.

### Topographic dissimilarity

#### Effect of stimulation orientation

TANOVA identified two intervals of significant dissimilarity: 10-59 ms and 66-106 ms. Within these intervals, local maxima of total explained variance (%EV) were found at 22, 43 and 81 ms post stimulus (Fig. 3a). These latencies, corresponding to AG-TEP components P25, N45 and N75, were then used to extract mean topographies for all compared datasets (Fig. 3b). Six post-hoc paired t-tests were performed at all three latencies but only comparisons at 43 and 81 ms were found significant. In both cases, a significant difference was observed between conditions of opposite coil position (*along* – *across*, t-maps shown in Fig. 3c), while no significance was found for the effect of current direction. At 43 ms (N45), the t-maps of all *along* – *across* t-tests showed very similar patterns, with a positivity centro-anterior on the stimulated hemisphere. The t-value distribution was less uniform at 81 ms (N75) but still featured in all four cases a centro-frontal ipsilateral positivity. The locations of t-map maxima and minima and the associated t-values are summarized in Table 1.

**Table 1.**
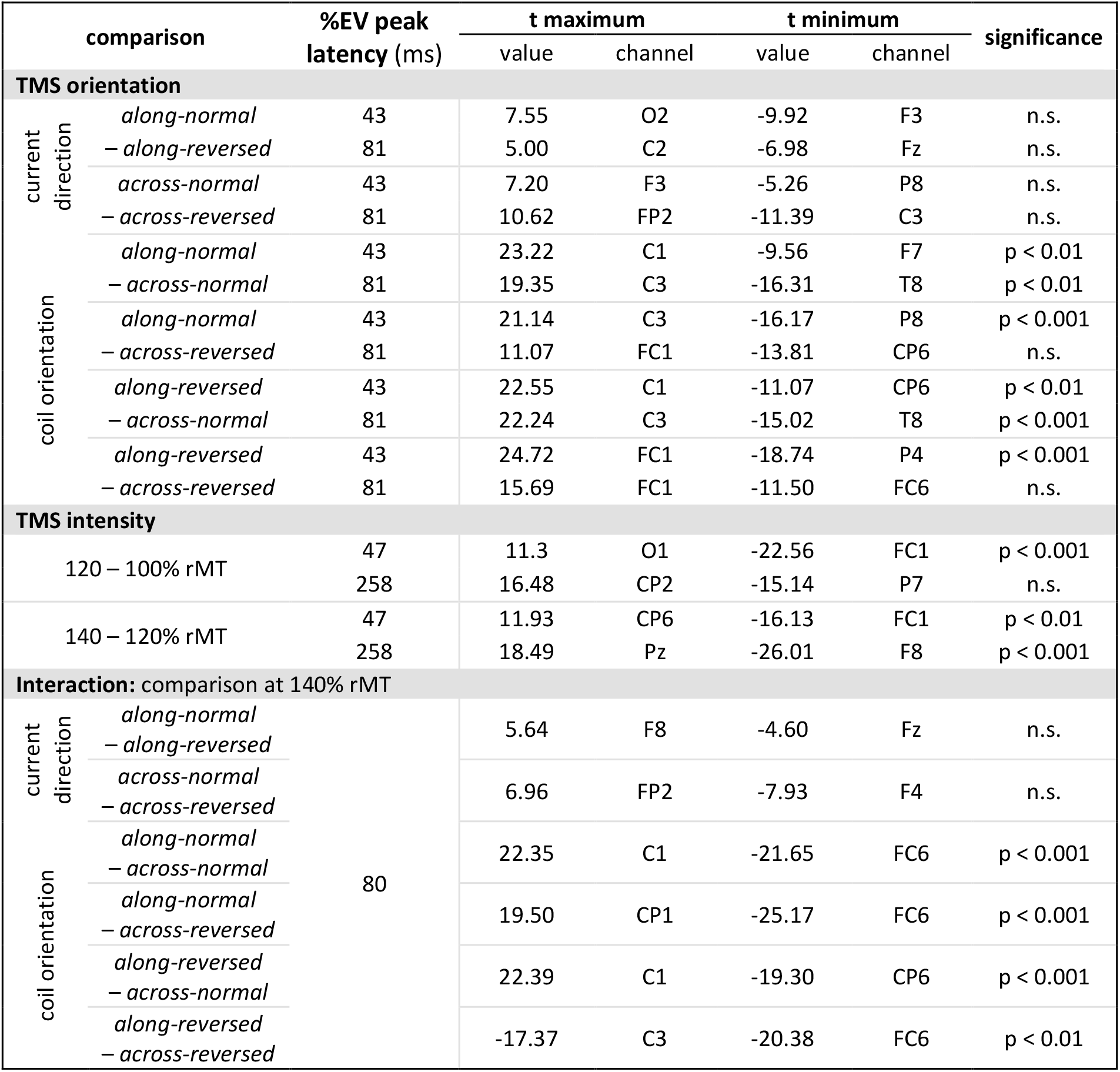
Post-hoc evaluation of topographic dissimilarity

**Fig. 3.**
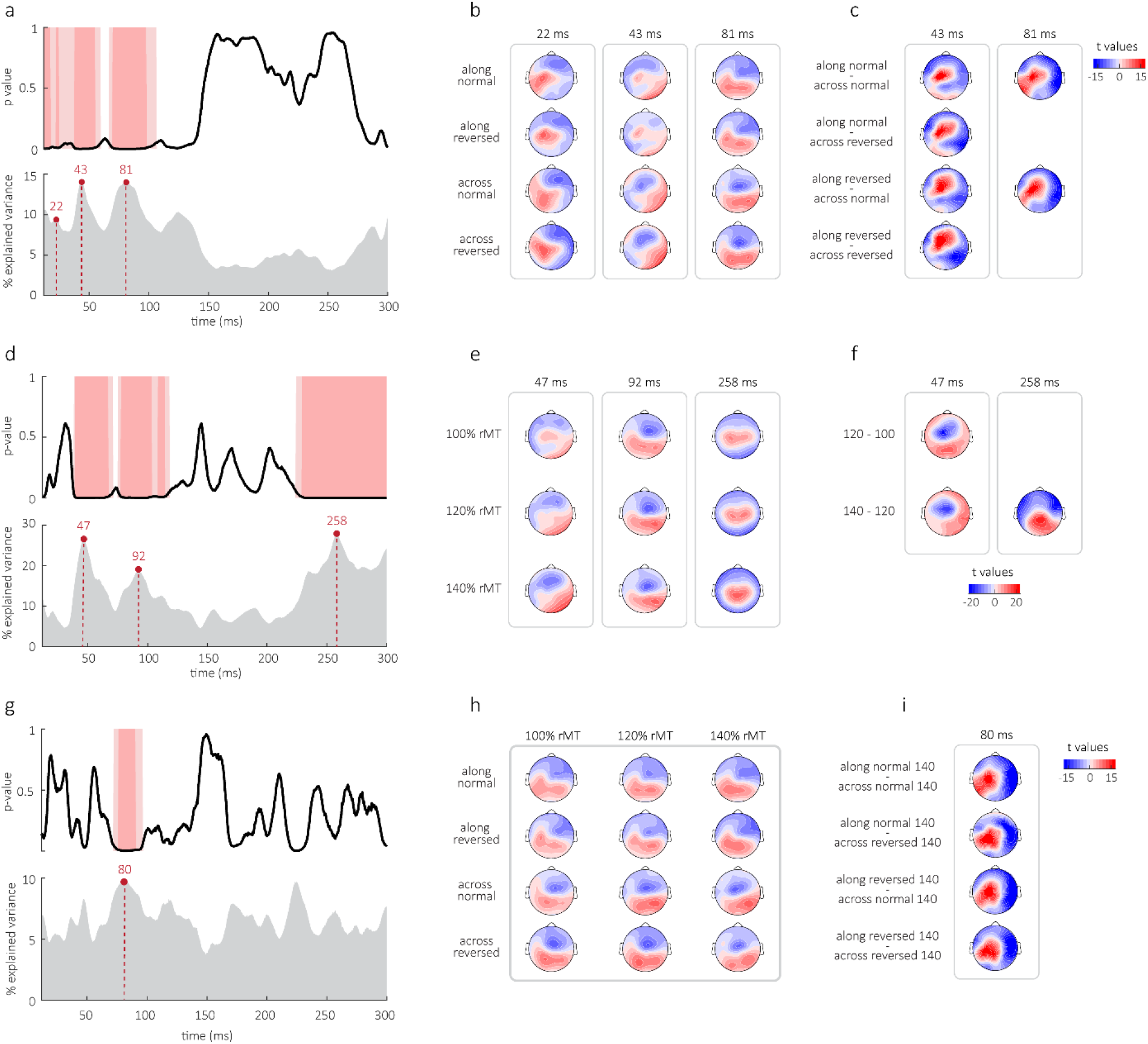
Analysis of topographic dissimilarity for different stimulation orientation (a, b, c), intensity (d, e, f) and the interaction of both factors (g, h, i). (a, d, g) Upper plots show p-values obtained by TANOVA in the analyzed time window (10 – 300 ms). Areas shaded in light pink correspond to intervals of p < 0.05, dark pink areas to intervals of p < 0.01. Bottom plots show percentage of explained variance attributed to given factor at each timepoint, peak latencies are marked by red dotted lines. **(b, e, h)** Mean topographies normalized to GFP that are associated to peak %EV latencies. **(c, f, i)** T-maps show the distribution of t-values obtained in post-hoc comparisons. Only the significant comparisons are shown.

#### Effect of stimulation intensity

TANOVA identified three intervals during which the topography significantly changed with the increasing stimulation intensity: 38-70 ms, 74.5-118 ms and 224-300 ms. Local maxima of %EV were found at 47, 92 and 258 ms, corresponding to AG TEPs components N45, N75 and P280 (Fig. 3d,e). At each %EV peak latency, post-hoc t-tests were performed to compare datasets evoked by two neighbouring stimulation intensities. At 47 ms (N45), the topography changed significantly between 100 and 120% rMT as well as between 120 and 140% rMT and both associated t-maps had a similar distribution pattern with the minimum localized frontally at the stimulated hemisphere and the maximum posteriorly at the opposite hemisphere. In addition, the topography was found significantly different when comparing datasets evoked by 120 and 140% rMT at 258 ms (P280), with the corresponding t-map showing an increase in positive values posterio-centrally (Fig. 3f, Table 1).

#### Interaction of stimulation orientation and intensity

One interval at 71.5-96 ms was retained by TANOVA as significant, with a local %EV maximum at 80 ms corresponding to the component N75 (Fig. 3g, h). Six post-hoc t-tests were used to compare all datasets at 100% rMT and at 140% rMT. While neither of the comparisons at 100% rMT proved significant, highly significant differences were found at 140% rMT when topographies with opposite coil position (*along* - *across*) were compared. All t-maps showed similar patterns, with positivity at the stimulated hemisphere (Fig. 3i, Table 1).

### Microstate analysis

The model (92.3% explained total variance) yielded six microstate classes (Fig. 4a) and attributed a prominent topography to each TEP component (Fig. 4b). Intervals of low TEP amplitude, particularly around 50 ms, were associated with rapidly switching short-lasting microstates, suggesting moments of topographic instability or transition (Fig. 4b,c). Analysis of temporal features was performed for all microstate classes, significant results were obtained for classes 2, 3 and 4.

**Fig. 4.**
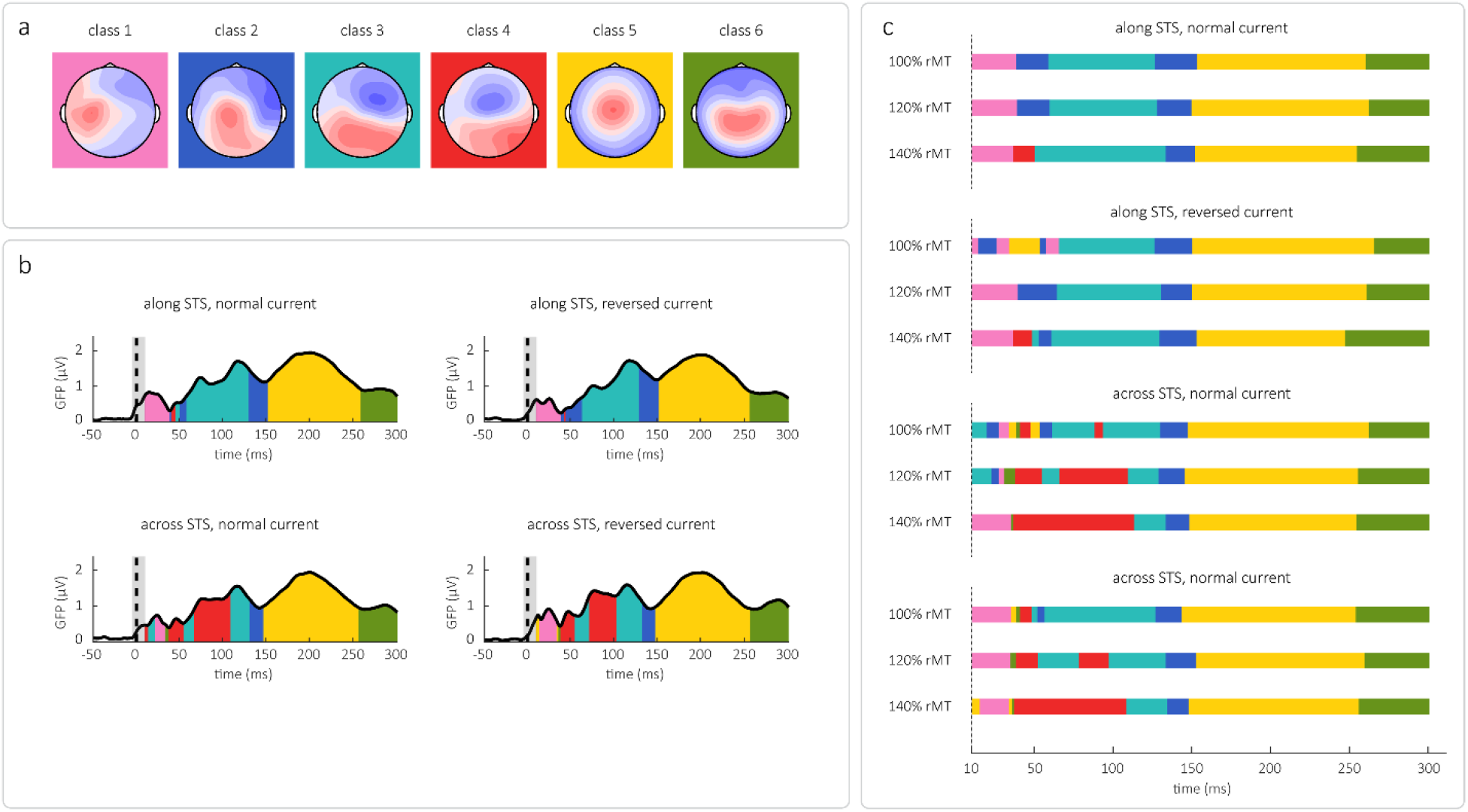
Microstate analysis. **(a)** 6 microstate class maps generated by the model with assigned colours. **(b)** GFP plots of TEPs averaged across tested intensities for each stimulation orientation are split into intervals corresponding to microstate classes to illustrate how microstates correspond to TEP components. **(c)** Microstate sequences of TEPs of all conditions; timepoints labelled with different microstate classes are colour-coded according to panel a.

Microstate class 2 appeared at the falling slope of component N100 and in most *along* datasets and in *across-100* datasets, it also appeared at the rising slope of N75. This was reflected in statistically significant effect of both orientation (p < 0.001) and intensity (p < 0.001) on its overall duration. We observed that class 2 lasted longer in *along-normal* and *along-reversed* datasets (on average 23 and 45 ms) than in *across-normal* and *across-reversed* datasets (15 and 16 ms). Its duration was reduced with increasing stimulation intensity (100% rMT – 40 ms; 120% rMT – 18.5 ms; 140% rMT – 17.5 ms) and this trend was more pronounced in *across* datasets (borderline significant).

Microstate class 3 represented the peak topography of both N75 and N100 in all datasets except for *across-normal-140* and *across-reversed-140*, where it was replaced at N75 latency by microstate class 4. While the occurrence of class 3 in *along* datasets changed only slightly with stimulation intensity, we saw its progressive suppression in *across* datasets as the intensity increased. This corresponds to the significant effect of orientation-intensity interaction on class 3 onset. Mean onset (ms) for 100 / 120 / 140% rMT: *along-normal* – 59.5 / 60 / 51; *along-reversed* – 66 / 65 / 49; *across-normal* – 62 / 55 / 114; *across-reversed* – 49 / 53 / 109.

The most pronounced differences between tested conditions were observed in temporal characteristics of microstate class 4. While it was missing in most *along* datasets, only appearing in a short interval in *along-normal-140* and *along-reversed-140* at latencies corresponding to N45, it was detectable in all *across* datasets where it also gained with the increasing stimulation intensity over the class 3 at the latency of N75. This is reflected in the significant effect of stimulation orientation and intensity, as well as their interaction, on the overall duration of class 4. Mean duration (ms) for 100 / 120 / 140% rMT: *along-normal* – 0 / 0 / 13; *along-reversed* – 0 / 0 / 12; *across-normal* – 12 / 61 / 77; *across-reversed* – 8 / 33 / 71. In addition, we found a significant effect of stimulation orientation on the class 4 offset. Mean offset (ms): *along-normal* – 46; *along-reversed* – 44; *across-normal* – 109; *across-reversed* – 103.

### Contribution of muscular contraction

During the study, a suspicion was raised that topographic differences observed between *along* and *across* AG-TEPs may be related to the characteristics of face muscle contractions directly evoked by TMS rather than differential activation of target brain networks. Therefore, an exploratory analysis was carried out to evaluate the contribution of this muscular artifact. A high-amplitude evoked potential peaking within 10 ms post stimulus and showing typical peripheral distribution (Fig. 5a) was recovered from untreated EEG data (TMS artifact was removed between -2 and 3 ms) and attributed to the muscular contraction. Its magnitude was quantified at the subject level as the average GFP_muscle_ at 3-10 ms post stimulus (only electrodes from the stimulated hemisphere were included; Fig. 5b). At this point, two additional subjects were identified as outliers based on their GFP_muscle_ values and removed (final n_muscle_ = 16). Group visualization revealed that *across* datasets were associated with a larger muscular artifact that showed a steeper increase with stimulation intensity as compared to *along* datasets, while the influence of current direction apparently less important (Fig. 5c). Mean GFP_muscle_ values are summarized in Table 2.

**Fig. 5.**
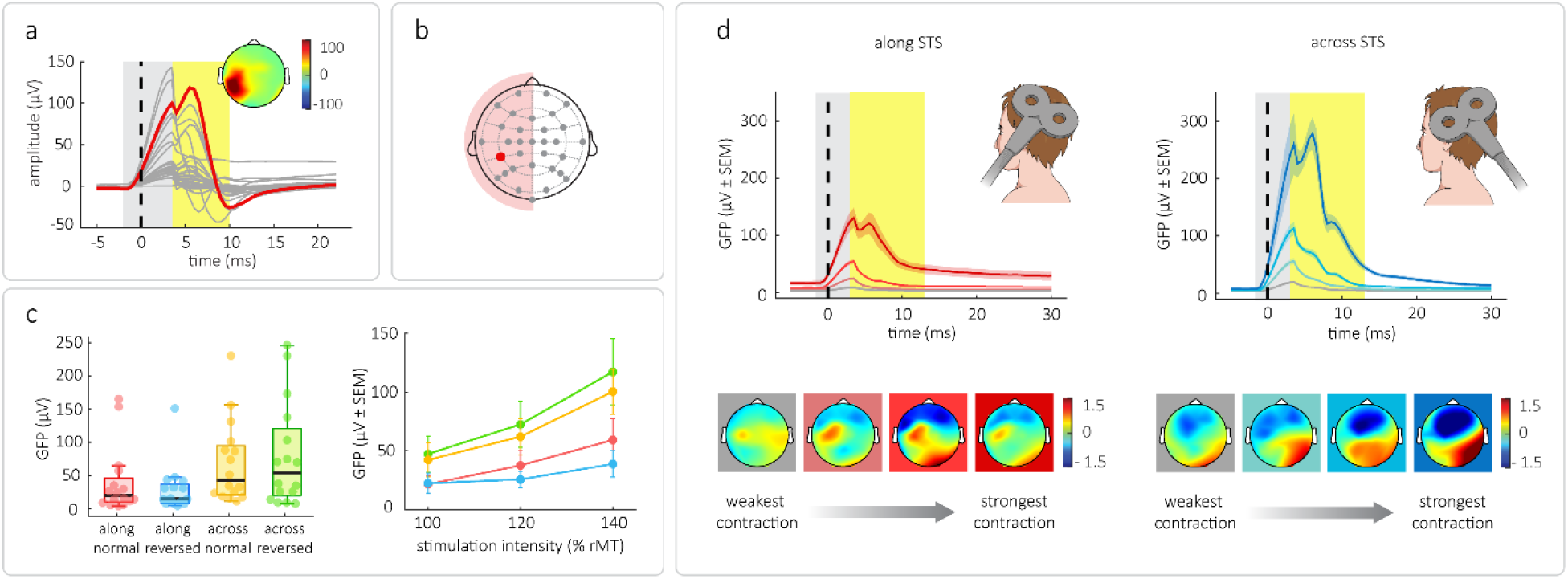
Quantification of evoked muscular contraction. **(a)** Average signal from all datasets showing the time-course and topography of the artifact attributed to muscular contraction; grey lines represent the signal of all electrodes, CP5 is highlighted in red. The time interval of TMS artifact interpolation is shaded in light grey, yellow area represents the time of interest over which the GFP_muscle_ was calculated. The topography was extracted at the latency of the CP5 peak (5 ms post stimulus). **(b)** EEG layout used in the study with the CP5 electrode highlighted in red marking the approximate site of stimulation. Only the signal from channels at the left (stimulated) hemisphere were used to calculate mean GFP_muscle_ used to quantify the strength of muscular twitch. **(c)** Mean group values of GFP_muscle_; red – along-normal, blue – along-reversed, yellow – across-normal, green – across-reversed. Left panel shows individual data with all intensities averaged; black bars mark median values, box whiskers connect lower and upper quartiles to nonoutlier maxima and minima. Right panel shows group average values (± SEM) for all datasets. **(d)** Increasing muscular artifact in along (red, upper panel) and across (blue, upper panel) datasets with the corresponding topography at 45 ms (lower panels). The interval of TMS artifact interpolation is shaded in light grey, yellow area represents the time of interest over which the GFP_muscle_ was calculated.

**Table 2.** Magnitude of TMS-evoked muscular contraction.

| dataset |  |  | dataset |  |  |
| --- | --- | --- | --- | --- | --- |
| orientation | intensity (%rMT) | average GFP<sub>muscle</sub> ($\mu\text{V} \pm \text{SEM}$) | orientation | intensity (%rMT) | average GFP<sub>muscle</sub> ($\mu\text{V} \pm \text{SEM}$) |
| along normal | 100 | $21.46 \pm 7.62$ | across normal | 100 | $42.27 \pm 14.48$ |
| | 120 | $37.55 \pm 12.52$ | | 120 | $62.01 \pm 15.37$ |
| | 140 | $59.18 \pm 18.52$ | | 140 | $100.45 \pm 19.30$ |
| along reversed | 100 | $22.28 \pm 8.70$ | across reversed | 100 | $47.29 \pm 15.43$ |
| | 120 | $25.62 \pm 6.98$ | | 120 | $72.44 \pm 19.78$ |
| | 140 | $38.76 \pm 11.25$ | | 140 | $117.24 \pm 28.23$ |

Microstate analysis indicated that the most remarkable difference between *along* and *across* datasets was the occurrence of the topographic pattern attributed to the microstate class 4 at the N45 and N75 latency. To investigate the association of the muscular artifact magnitude and this specific topography, all data were grouped by coil position, sorted, and split into four categories of increasing GFP_muscle_ (Fig. 5d, upper panels). TEP topographies were extracted at 45 ms from processed datasets averaged within each category. We observed that in both *along* and *across* datasets, larger muscular artifacts were associated with more prominent class 4 topography (Fig. 5d, lower panels). Its appearance was less pronounced in *along* datasets, which corresponded to a comparatively weaker average muscular artifact.

## Discussion

In the present work, we characterized the spatiotemporal profile of AG-TEPs and showed that its topographic features depend on TMS coil orientation and stimulation intensity. Specifically, when investigating the effect of coil orientation, we found topographic dissimilarity across tested conditions at latencies corresponding to components P25, N45 and N75, indicating that different cortical sources were active or that the same sources contributed differently to the global activity. However, it appears that this variability cannot be solely attributed to differences in activity of the stimulated area/.

One major concern when interpreting TEPs is the inevitable contamination by peripherally evoked sensory potentials (PEPs) (Biabani, Fornito, Mutanen, Morrow, & Rogasch, 2019; Conde et al., 2019). The magnetic field and coil vibration activate a range of receptors in the soft tissues of the head, thereby triggering somatosensory-evoked brain potentials (SEPs), while the clicking sound of TMS produces auditory-evoked potentials. By comparing active TMS with sham stimulation, multiple studies showed that PEP contribution to TEPs increases with time and must be considered as soon as 50-60 ms post stimulus (Biabani et al., 2019; Chowdhury et al., 2022; Freedberg et al., 2020; Pedro Caldana Gordon, Desideri, Belardinelli, Zrenner, & Ziemann, 2018; Pedro C Gordon et al., 2021; Poorganji et al., 2021). Similarly, in our previous study (Sulcova et al., 2022b), we showed that topographic features of TEPs evoked by stimulation of different cortical regions become progressively more similar and components peaking later than approx. 80 ms likely mostly reflect target-unspecific activity attributed to PEPs. Therefore, P25 and N45 should be the most relevant components of AG-TEPs for the assessment of the functional state of the AG.

Several arguments support the claim that P25 likely reflects neural activity directly evoked within the AG and connected areas. As we previously showed, the topography of this component is highly target specific (AG-TEPs were recorded with the coil placed across STS) (Sulcova et al., 2022b). Here, we observed significant topographic dissimilarity of P25 of AG TEPs evoked by different stimulation orientation, yet the patterns were similar enough to be pooled into a single microstate. This subtle variability could be explained by a differential engagement of local neuronal sub-populations depending on their sensitivity to different features of the induced electric field. Moreover, another recent study succeeded to locate P25 cortical generators in the stimulated AG as well as the ipsilateral ventral medial prefrontal cortex (vmPFC) (Ozdemir et al., 2020), an area also belonging to the DMN (Raichle et al., 2001). Therefore, in the future, AG-TEP component P25 may serve as a non-invasive biomarker of the functional properties of the AG and the networks it integrates.

By contrast, our data suggest that the later AG-TEP components might be prone to contamination by somatosensory-evoked neural activity, and more specifically, by the brain response to the TMS-evoked twitch of head muscles. When the TMS coil is placed over a head muscle, the stimulation triggers its contraction (Machetanz et al., 1994) which, in turn, activates mechano-sensitive somatosensory afferents. Because coil placement determines the character and extent of muscle activation, distinct coil orientations are likely associated with differences in the strength of muscular activation and somatosensory feedback, leading to variable degrees of TEP contamination by somatosensory-evoked cortical activity. These differences might be especially pronounced when stimulating targets with a lateral projection on the scalp, such as the AG.

Increasing stimulation intensity also evokes a stronger muscular contraction that elicits a stronger associated SEP, and this increase in SEP amplitude presumably follows different dynamics than the TMS intensity-dependent increase in directly evoked cortical activity. Because the head muscles are engaged by the strong electric field in the TMS coil proximity, even a small increase of stimulation intensity likely leads to a considerable augmentation of muscular contraction that translates into a steep increase of SEP amplitude. On the other hand, direct TMS activation of cortical neurons is triggered by much weaker electric fields and the final strength of resulting brain response is dampened by parallel co-activation of different neural components. It can be therefore expected that the amplitude of neural activity directly evoked within the stimulated network grows slower than the amplitude of SEPs and, by consequence, the TEP signal is progressively dominated by SEPs as the intensity increases.

Here, we observed that the TMS-evoked muscular contraction was stronger when stimulation was delivered *across* as compared to *along* the STS and augmented with increasing stimulation intensity. Although TMS stimulation directed across the STS should lead to optimal activation of AG neurons in the sulcal wall, this coil placement visibly engages head muscles, such as *m. temporalis*, and should be therefore avoided. Because all four coil orientations yielded a clear (if not completely identical) P25 and therefore presumably captured the activity within the stimulated network, the coil orientation may be in future studies adjusted at the individual level to minimize the interaction with head muscles. This could be achieved for example by real-time visualization of the average TEP signal, a recently developed method that allows to minimize muscular artifacts while maximizing the early, most relevant TEP components (Casarotto et al., 2022).

Importantly, our results showed that the effects of TMS coil orientation and stimulation intensity on the N45/N75 topography followed the effects on TMS-evoked muscular activity. The microstate analysis revealed that these AG-TEP components elicited by stimulation *across* the STS was dominated by a bipolar topographic pattern (microstate class 4) centred over the contralateral hemisphere, whose occurrence and duration increased with stimulation intensity. Interestingly, this topography bears a striking resemblance to that of mid-latency SEPs evoked by electrical stimulation of mixed peripheral nerves of the face accompanied by a muscle twitch (Buchner et al., 1994; Miller, Longo, & Saygin, 2019; Restuccia et al., 2002). Independently of coil position, this pattern became more pronounced as the magnitude of the muscular artifact increased, which would be consistent with it reflecting the somatosensory response to the muscle contraction. Furthermore, the microstate in question tended to span both N45 and N75 when the stimulation was delivered with the highest intensity *across* the ST (thus leading to the strongest muscular contractions). Therefore, we propose that even in TEPs elicited with lower intensities, where the occurrence of this microstate was not continuous, it may represent the same SEP signal overlaying the ongoing TMS-evoked neural activity and temporarily dominating at both latencies.

Lastly, considering the influence of individual anatomy on muscular contractions, variable occurrence of muscular activation could explain the inter-subject topographic inconsistency of TEP components peaking around 45 ms. Similar inconsistency was previously also described for motor cortex TEPs (Sulcova et al., 2022b), showing that this period of excessive inter-subject variability is surprisingly conserved in time and across conditions, as could be expected in case of variable SEP contribution. These findings indicate that any TEP changes occurring as soon as approx. 45 ms should be interpreted with caution, especially when simultaneous changes in evoked muscular activity are observed.

## Conclusion

In summary, we show that topographic features of early AG-TEPs are dependent on TMS stimulation parameters, thus reflecting differential engagement of underlying brain sources. We conclude that the earliest component P25 of AG TEPs likely corresponds to brain activity evoked by direct activation of the AG and as such could serve as a biomarker of excitability within the AG and the connected DMN. In contrast, the subsequent components N45 and N75 seem to be influenced by the presence of TMS-evoked peripheral muscle activity and should be interpreted with caution. As all tested TMS coil positions efficiently activate the AG, the coil placement should be adjusted to minimize muscular contractions when recording AG-TEPs.

## Acknowledgements

Presented work was supported by the Fund for Scientific Research (F.R.S.–FNRS; Wallonia-Brussels Federation) of which D. Sulcova was a research fellow.

