## Supplementary materials for "Exploring the properties of the left angular gyrus using TMS-evoked potentials" (from another version of this preprint: version 1, 2022-11-15)

### SUPPLEMENTARY MATERIALS: Influence of stimulation parameters on AG TEP magnitude

#### Material and methods

The analysis of topography provides valuable information about the qualitative changes in the EEG signal, but since it operates with normalized data, it disregards the quantitative changes in the EEG amplitude. For this reason, we also performed all planned comparisons considering the magnitude of AG TEPs, which was expressed as GFP and evaluated using a point-by-point randomization-based test implemented in Ragu. 5000 permutations were run to obtain an empirical distribution of values true under the null hypothesis and p value was extracted as a function of time. The clusters of significant timepoints were retained only when surviving the 'Global Duration Statistics' correction.

#### Results

While we did not see any effect of stimulation orientation on the GFP, the effect of stimulation intensity was expectedly found significant over the whole considered time window. When considering the interaction of both factors, an interval of significance was identified at 17 – 56.5 ms post stimulus, which includes the earliest TEP components P25 and N45 (Fig. S1a). Remarkably, a sharp local maximum of %EV was localized at 42 ms post stimulus (Fig. S1b) and the time interval covering N45 (30.5 – 55 ms) was characterized by very low p values ($p < 0.01$).

To further explore the effect of interaction at N45, GFP was averaged across the interval of $p < 0.01$ at the level of individual subjects and mean values were extracted for each tested condition (Fig. S1c). We observed that when the stimulation intensity of 100% rMT was used, response strength was greater in *along* datasets than in *across* datasets, While the GFP values generally increased with the stimulation intensity, this increase was much steeper in *across* datasets, leading to larger GFP values at 140% rMT in *across* than *along* datasets. The trend was further accentuated when visualized as the GFP change with increasing intensity (Fig. S1d).

Average values (µV ± SEM) obtained for each condition (100 / 120 / 140% rMT): *along-normal* – 1.14 ± 0.11 / 1.26 ± 0.11 / 1.47 ± 0.14; *along-reversed* – 1.08 ± 0.08 / 1.27 ± 0.12 / 1.32 ± 0.10; *across-normal* – 0.94 ± 0.06 / 1.26 ± 0.11 / 2.03 ± 0.22; *across-reversed* – 0.99 ± 0.06 / 1.41 ± 0.12 / 1.98 ± 0.19.

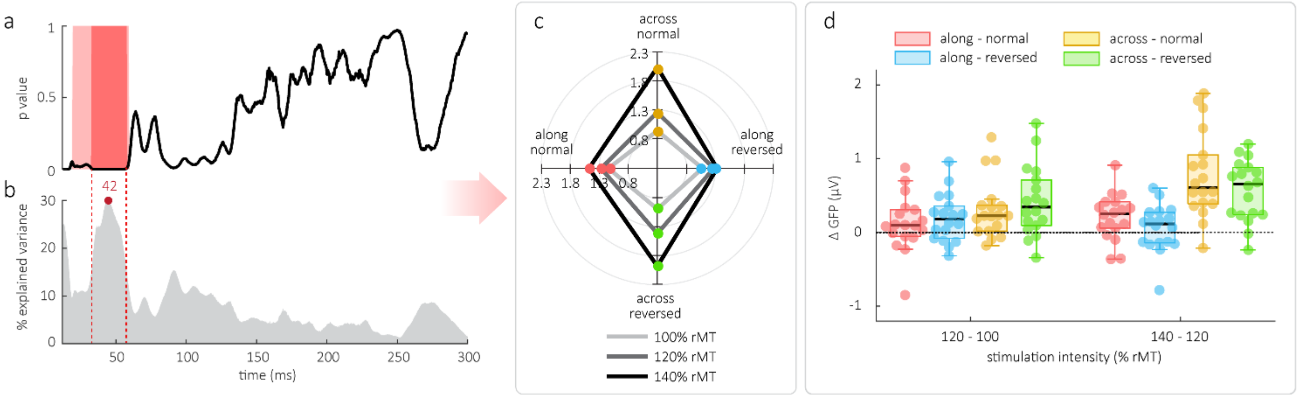


***Fig. S1 - Analysis of TEP amplitude.*** (a) P-values obtained by permutation-based ANOVA across the whole analysed time window (10 – 300 ms) for the interaction between factors *stimulation orientation* and *stimulation intensity*. Area shaded in light pink corresponds to the interval of $p < 0.05$, dark pink area to the interval of $p < 0.01$. (b) Percentage of explained variance attributed to each timepoint. Red dotted lines indicate the time interval that was used to extract the mean GFP; the peak latency is denoted by a red dot. (c) Radar plot showing the mean GFP averaged across the interval of interest in all compared conditions. The axes represent levels of stimulation orientation; levels of stimulation intensity are marked by full lines (see legend). (d) The change in mean GFP with the increasing stimulation intensity. The change was calculated at the individual level as a simple subtraction of the GFP at lower intensity from the GFP at the higher intensity. Levels of stimulation orientation are distinguished by colour (see legend), black bars mark median values, box whiskers connect lower and upper quartiles to nonoutlier maxima and minima.
